# Resolving the neural processes involved in affective self-other distinction – an fMRI study

**DOI:** 10.64898/2026.11.17.764825

**Authors:** Tatiana Goregliad Fjaellingsdal, Konstantin Döhr, Emily Fitzgibbon, Martin Göttlich, Ulrike M. Krämer

## Abstract

During interactions, we need to distinguish our own emotions from our empathic response to the emotions of others. This ability of affective self-other distinction (SOD) is particularly challenged when direct cues about others’ emotions, such as emotional facial expressions, activate shared representations of affective experience. Employing a novel affective SOD paradigm alongside fMRI, we examined the neural processes associated with affective SOD. In this paradigm, dynamic emotional facial expressions that are either congruent or incongruent with the participant’s own emotional state are presented, and the participant must judge their own or the other person’s experience. Insufficient SOD can lead to an emotional egocentric or altercentric bias, depending on whether one’s own or another’s emotions are being evaluated. Previous studies have shown that these biases are modulated by self-salience, i.e., the extent to which one prioritizes information about oneself. Our study therefore aimed to clarify the distinct and joint neural correlates of affective SOD and self-salience. Data from 24 healthy young participants were analyzed. The results showed that incongruence between one’s own emotions and those of others activated the right supramarginal gyrus (rSMG) and the anterior insula. This extends previous studies of SOD in which participants had to infer others’ emotions indirectly based on emotion-eliciting stimuli. Our findings suggest a more general, modality-independent role for the rSMG in SOD. Furthermore, self-salience was associated with increased activity in parts of the default mode network and the rSMG, suggesting that SOD and self-salience rely on common neurocognitive processes.

## Introduction

When we interact with other people, we have to represent the other’s mental state in order to understand and predict their behavior. We do so by inferring their mental states from the context (“theory of mind” route) or by sharing their experience based on cues such as facial expressions (“experience sharing” route) (Frith & Frith, 2006; Lieberman, 2007). What happens though, when our own mental state, especially emotional state, is inconsistent with the state observed in the other? How do we keep track of conflicting representations of our own and the other’s affective state? The ability to deal with this challenge has been called affective self-other distinction (affective SOD) (de Guzman et al., 2016; Eddy, 2018; Lamm et al., 2016; Silani et al., 2013; Steinbeis, 2016), and difficulties in this ability are thought to be characteristic of certain psychiatric conditions, e.g., Borderline Personality Disorder (De Meulemeester et al., 2021).

Affective self-other distinction becomes most relevant in situations of incongruence between the observed emotional states in others and one’s own emotional state. An emotional egocentric bias (EEB) reflects the tendency to project one’s own emotional state onto the other and perceive the other’s emotion as more similar to one’s own state than it actually is. An altercentric bias (EAB) on the other hand occurs when the other’s emotional state affects me in a way that my perception of my own emotion becomes biased towards the other’s state. Additionally, the perception of others’ emotions might be biased because we generally tend to prioritize information pertaining to ourselves compared to information pertaining to others, a phenomenon called self-salience. Studying interindividual differences in EEB and EAB requires accounting for self-salience (Bukowski et al., 2021; Goregliad Fjaellingsdal et al., 2025), yet no fMRI study on affective SOD has done so.

In an influential study on the neural basis of affective SOD (Silani et al., 2013), the authors showed that the right supramarginal gyrus (SMG) is activated specifically when evaluating the other’s affective states which are incongruent to one’s own state, and that inhibition of the rSMG using offline TMS led to an increased EEB. Two more imaging studies found rSMG activation for incongruent compared to congruent affective states, but this effect was not specific for the EEB, but was also found for the EAB (Bukowski et al., 2020; Riva et al., 2016). Overall, previous studies speak for an important role of the rSMG for affective SOD, but its activity and interaction within larger neural networks seems to depend on the task setting and modality (Hartmann et al., 2022; Steinbeis et al., 2015).

What these previous affective SOD paradigms have in common is that the affective state of the other person has to be inferred indirectly from the eliciting situation or stimulus. We reasoned that the direct perception of cues about the other’s affective states (e.g., facial expressions) should activate the “experience sharing” route and should challenge affective SOD more (Goregliad Fjaellingsdal et al., 2025). In the “Food Emotional Egocentricity Paradigm” (FEEB), participants have to judge their own or another person’s emotional response to food pictures based on brief videos of another person bearing a happy or disgusted facial expression. Here, we used this paradigm together with fMRI to further elucidate the neural basis of affective SOD. We asked whether the same rSMG region is recruited when conflicting representations are activated through the “experience sharing” route and what the neural correlates of affective self-salience are.

Based on the suggested modality- and domain-general function in affective self-other distinction, we hypothesized that the rSMG should be activated in cases of incongruent emotional states. Given their roles in experience sharing, we expected the anterior cingulate cortex (ACC) and anterior insula (aIns) to be activated in conditions of affective incongruence. We hypothesized that evaluating one’s own affective state compared to that of others would activate default mode network regions (Mars et al., 2012; Schurz et al., 2020).

## Methods

### Participants

Forty-five self-reportedly healthy participants (29 female, 16 male) with no history of psychiatric or neurological diseases took part in this study. Participants had a mean age of 23.4 ± 4.1 years, ranging from 18 to 40 years. Recruitment of participants took place via university mailing lists and participation was compensated (€10 per hour or course credits). The study was conducted following the Declaration of Helsinki principles and was approved by the ethics committee of the University of Lübeck (ethics vote number 2023-288). We aimed for a sample size of n = 40, which was based on previous fMRI studies on SOD (Bukowski et al., 2020; Silani et al., 2013) and expecting an exclusion rate of about 20%. As our final exclusion rate was higher, we increased our sample size. Of all tested participants, one had to be excluded because of artifacts, one because of anatomical issues, and one because of software problems. Another five had to be excluded because of excessive head motion. From the remaining 37 participants, 13 participants were excluded because they met either or both of the FEEB exclusion criteria (see below), resulting in 24 participants (18 female and 6 male) who remained for the analysis. The mean age of the final sample was 22.6 ± 3.0 and ranged from 19 to 34 years.

### Paradigm

The Food Emotional Egocentricity Bias (FEEB) (Goregliad Fjaellingsdal et al., 2025) paradigm was used to study affective SOD. The paradigm simulates the experience in a restaurant setting, where food is served (i.e., a picture of the food is shown) and the participant then looks up to see the emotional reaction of the other person (i.e., a video of a person displaying happiness or disgust) at the table. The FEEB paradigm has two parts: The first part consists of a self-paced *prerating*, during which each food picture (n=20) is presented in a randomized fashion and has to be rated by the participant on a 10-point scale of -5 (very aversive) to +5 (very appetitive). The subjective rating is used to classify the participant’s preference towards that food item. During the second part, the *main task*, participants have to rate their own or the other person’s emotion towards the presented food (on the same 10-point scale). Each individual trial consists of a fixation cross (2 – 7 s), the food picture (2 s), the video of the other’s emotional facial expression in the middle of the screen together with the food picture in the lower part of the screen (3 to 4 s), another fixation cross (1.5 – 2.5 s), the presentation of a colored circle which indicates the target to be rated (2 s), and finally the rating scale (presented until a rating was given, time-out after 5 s). To keep the trial length constant across trials, the presentation time of the fixation cross is set to be 2 s plus the remaining time of the prior trial’s maximal rating duration of 5 s (see Figure 1). Participants performed two practice trials prior to the measurement starting to get accustomed to the structure of the main task. The colors indicating self or other target were counterbalanced between participants.

**Figure 1.**
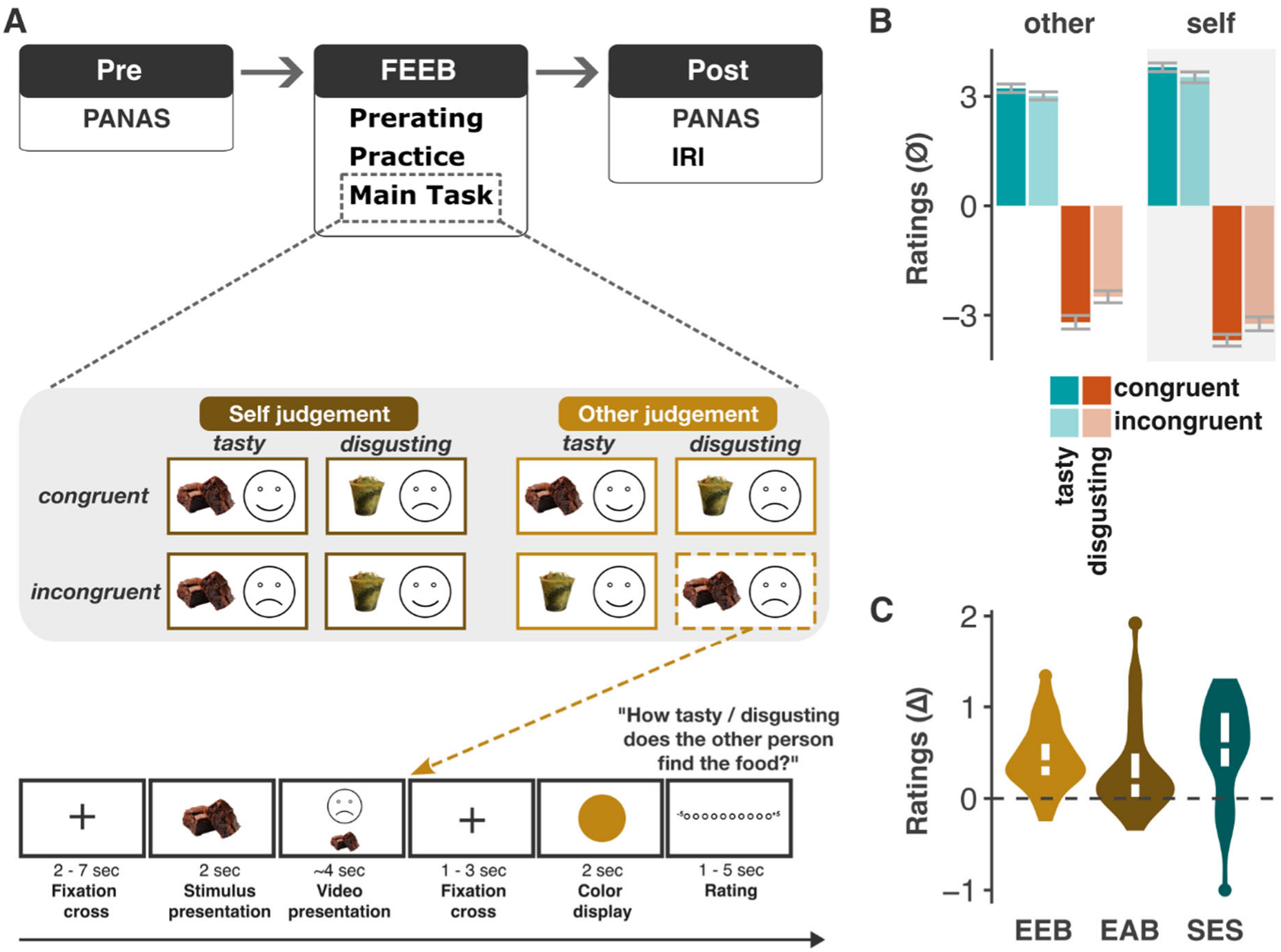
Session overview and behavioral results. **(A)** An overview of the session including questionnaires and the FEEB. Each trial of the main task can be categorized by the target (self / other), the valence (tasty / disgusting), and the congruency (congruent / incongruent). At the bottom is an example for an incongruent trial where the other person’s disgusted response has to be evaluated. Note that the “smiley” is only an illustration of the actual stimuli, which were short videos of persons showing a happy or disgusted face. Video stimuli were taken from the Jerusalem Facial Emotion Expression Set (JEFEE) (Yitzhak et al., 2017). **(B)** Mean ratings and standard errors during the main task on a scale from -5 to +5 **(C)** Distribution of EEB (golden), EAB (dark golden), and SES (teal).

The FEEB paradigm has a 2 x 2 x 2 design with the factors Congruency (congruent vs incongruent), Target (self vs other) and Valence (tasty vs disgusting) (Figure 1). The factor Congruency pertains to the congruency between the participant’s emotional response to the food (based on the pre-rating) and the depicted emotional expression in the video. The factor Target refers to whether the participant has to evaluate their own or the other’s emotion. Finally, the factor Valence refers to the valence of the food for the to-be-evaluated person in this trial. When the participant had to rate their own emotion (self-rating trials), stimuli were classified as having positive valence (tasty) when preratings were positive (≥1), and negative valence (disgusting) when preratings were negative (≤-1). For other-rating trials, valence was based on the emotional expression displayed in the video stimulus, with happiness indicating a tasty valence and disgust indicating a disgusting valence. The presentation of the trials was pseudorandomized to avoid having more than three consecutive trials of the same valence in the video or the same target to be rated, or more than two consecutive presentations of the same food stimulus.

### Materials and questionnaires

The materials for the FEEB paradigm consisted of food pictures and video stimuli. The food pictures were taken from the Food Pics Database (Blechert et al., 2019) and Food-475 Database (Ciocca et al., 2018). The video stimuli were adapted from the Jerusalem Facial Emotion Expression Set (JEFEE) (Yitzhak et al., 2017) and shortened to 3-4 s long displays of smiling faces and disgusted faces with a short neutral face beforehand. The paradigm was programmed in Matlab (MathWorks Inc) using Psychophysics Toolbox (Brainard, 1997; Kleiner et al., 2007; Pelli).

The questionnaires included a pencil-based lab-internal MRI safety questionnaire, and via Labvanced (www.labvanced.com), we administered the following questionnaires: Positive Affect and Negative Affect Schedule (PANAS) in German (Janke & Glöckner-Rist, 2012), Interpersonal Reactivity Index (IRI) in German (Davis, 1983; Paulus, 2009), a customized evaluation questionnaire (e.g., ‘How well could you rate the experience of the other person?’/’How well could you rate your own experience?’, both on a scale from 1 = very poorly to 10 =very well), and a query of the emotion displayed in each of the presented videos (multiple choice selection of an emotion after being displayed the video alone).

### Procedure

On the day of the measurement, participants first signed the informed consent. A screening questionnaire for MRI inclusion criteria and the PANAS were administered before the measurement. Thereafter, the MRI session (see MRI data acquisition for detailed protocol) started with the FEEB paradigm. After the MRI session, participants filled out the PANAS, the IRI, and an evaluation questionnaire (Figure 1). They also rated for each of the videos which emotion was displayed.

### MRI data acquisition

Structural and functional MR imaging was performed at the CBBM Core Facility Magnetic Resonance Imaging using a 3-T Siemens Magnetom Skyra MRI scanner equipped with a 64-channel head-coil. Functional images were acquired applying a single-shot gradient-recalled echo-planar imaging (GRE-EPI) sequence sensitive to blood oxygen level dependent (BOLD) contrast (Time to repeat = 980 ms; echo time = 30 ms; flip angle = 65°; in-plane resolution 3×3 mm^2^; 204×204 mm^2^ field of view; 56 transversal slices; 3 mm slice thickness; simultaneous multi-slice factor 4; 802 volumes were recorded). Additionally, structural images of the whole brain using a 3D T1-weighted MP-RAGE sequence were acquired (TR=1900 ms; TE=2.44 ms; TI=900 ms; flip angle 9°; 1×1×1 mm^3^ resolution; 192×256×256 mm^3^ field of view; acquisition time 4.5 minutes).

### fMRI preprocessing

Functional MRI data were preprocessed using the Statistical Parametric Mapping software package (SPM12; Wellcome Centre for Human Neuroimaging, University College London, UK) implemented in MATLAB (The MathWorks, Inc., Natick, MA, USA). Slice-timing correction was applied to the functional images to account for temporal differences in the acquisition of individual slices using the actual slice acquisition times during SMS acquisition. Subsequently, the functional images were realigned to correct for head motion using SPM12’s rigid-body realignment procedure. A mean functional image was generated during the realignment procedure. For anatomical-functional alignment, the participant’s high-resolution T1-weighted anatomical image was co-registered to the mean realigned functional image. The T1-weighted anatomical image was subsequently segmented using the tissue segmentation procedure implemented in SPM12. The deformation field obtained from the segmentation of the T1-weighted image was then used for spatial transformation of the functional images from native anatomical space to standard Montreal Neurological Institute (MNI) space. Finally, the normalized functional images were spatially smoothed using a three-dimensional Gaussian kernel with a full width at half maximum of 8×8×8 mm^3^. The resulting preprocessed functional images were subsequently used for first-level statistical analysis.

### Behavioral analysis

Behavioral analyses were conducted using R Statistical Software (Version 4.4.2; R Core Team, 2024). Participants were excluded from subsequent analyses if they met any of the following criteria in the main task: (i) fewer than seven trials in at least one of the eight experimental conditions, or (ii) ratings exceeding predefined upper or lower thresholds in more than five trials. To determine these thresholds, ratings were averaged separately for each valence category across all conditions and participants. Interquartile ranges (IQRs) and corresponding quartiles (0.25 and 0.75) were calculated, and upper and lower bounds were defined as the respective quartile plus or minus 1.5 times the IQR. The number of trials in which each participant’s ratings fell outside these bounds was then computed.

Statistical analysis was performed on the averaged values for each participant and each condition. Data were analyzed using a repeated-measures ANOVA including three within-subject factors with two levels each: Congruency (congruent, incongruent), Target (self, other), and Valence (appetitive, aversive). Due to the structure of the data (valence negative and positive), the absolute value of negative ratings was used for the analysis. When significant interaction effects were observed, post hoc comparisons were conducted. Effect sizes are reported using partial eta squared (ηp^2^).

Emotional bias indices were operationalized as egocentric and altercentric biases (Goregliad Fjaellingsdal et al., 2025). The egocentric bias was computed as [(−1 × Δ1 + Δ3) / 2], whereas the altercentric bias was calculated as [(−1 × Δ2 + Δ4) / 2]. Here, Δ1 represents the difference between incongruent and congruent trials for pleasant ratings of the other target; Δ2 represents the difference between incongruent and congruent trials for pleasant self-ratings; Δ3 represents the difference between incongruent and congruent trials for unpleasant ratings of the other target; and Δ4 represents the difference between incongruent and congruent trials for unpleasant self-ratings. For the calculation of self-salience, the difference was computed between self- and other-ratings on congruent trials, which was averaged across positive and negative valence conditions.

For the PANAS, individual positive affect scores were obtained by summing all positive affect items, and negative affect scores were derived by summing all negative affect items (see table S1 for means and SD). Associations between PANAS scores and emotional bias measures were examined using Spearman rank-order correlations, with full results provided in the supplementary materials (see table S2). For the IRI, individual subscale scores for perspective taking (PT) and personal distress (PD) were calculated by summing the respective items for each subscale. Associations between IRI measures and emotional bias measures were examined using Spearman rank-order correlations, with full results provided in the supplementary materials (see table S2).

### fMRI data analysis

Subject and group level analyses were performed using SPM12 and the Marsbar toolbox (version 0.44) was used to extract mean contrast values in predefined regions of interest (ROI). For the first level analysis (subject level), we used the general linear model (GLM) approach implemented in SPM12. The following ten regressors were used: eight regressors of interest for the rating phase, separately for the eight conditions (0-5 seconds, depending on the participants reaction times), and two regressors of no interest (one regressor for the picture presentation (∼2s) and one for the video presentation (∼4s). The resulting model included a high-pass filter with a cut-off period of 128 s, and temporal autocorrelations were modelled using a first-order autoregressive model. We generated contrast maps for incongruent vs. congruent trials for other-ratings (SOD_other_), incongruent vs. congruent trials for self-ratings (SOD_self_), incongruent vs. congruent trials across self- and other-trials (SOD), the double difference between incongruent vs. congruent in other- and self-rating trials (SOD_other-self_; see Silani et al., 2013), and for congruent trials in self-ratings vs. congruent trials in other-ratings (self-salience). We set up an additional GLM model with separate four regressors for the video presentation in the four conditions of positive and negative valence in congruent and incongruent trials (e.g., happy video with congruent picture; ∼4s). We then created an additional contrast map for video presentation in incongruent vs congruent trials (ConVid). For all contrasts, we averaged across positive and negative valence.

For whole brain analysis on level 2 (group level), we first used simple t-tests to quantify the six different effects on brain activity (i.e., SOD, SOD_other_, SOD_self_, SOD_other-self_, self-salience, and ConVid). Afterwards, we used multiple regressions analyses to test the correlation between the three behavioral bias scores (EAB, EEB, and SES) and the respective contrast (e.g., EEB with SOD_other_). For all whole brain analyses, we used family-wise-error correction (FEW) at the cluster level (*p* < .05). The initial threshold was *p*_*thr*_ < .001. We used the automatic anatomical labelling atlas 3 (AAL3) to localize our local maxima (Rolls et al., 2020).

Our regions of interest (ROI) analysis focused on the right supramarginal gyrus (rSMG), which has been reported to be related to SOD/EEB (Bukowski et al., 2020; Silani et al., 2013). We used a 6mm sphere around the coordinates x = 65, y = -37, z = 33 in MNI space. We used the following contrasts for the ROI analysis: SOD, SOD_other_, SOD_self_, SOD_other-self_, self-salience, and ConVid.

## Results

### Behavioral results

A repeated measures ANOVA with three within-subject factors (congruency, target, and valence) yielded the following results (Figure 1): The participants gave more extreme ratings (more positive/negative) in congruent trials compared to incongruent trials (main effect of congruency: *F*(1,23) = 28.68, *p* < .001, *η*_*p*_^*2*^ = 0.555). Additionally, they rated their own emotional response as stronger than the other person’s emotion (main effect of target: *F*(1,23) = 18.90, *p* < .001, *η*_*p*_^*2*^ = 0.451), and they gave more extreme pleasant than unpleasant ratings (main effect of valence: *F*(1,23) = 4.82, *p* = .039, *η*_*p*_^*2*^ = 0.173). We also observed that the difference between congruent and incongruent ratings was higher for unpleasant stimuli than for pleasant ones (interaction effect of congruency and valence: *F*(1,23) = 8.53, *p* = .008, *η*_*p*_^*2*^ = 0.271). There was, however, no difference in congruency effects between ratings for oneself compared to rating pertaining to the other person (interaction effect of congruency and target: *F*(1,23) = 0.61, *p* = .44). This means that EEB and EAB did not differ from one another. We then calculated EEB, EAB, and SES (see Methods). Across all participants, all three bias scores differed significantly from zero (all *p* < .01), which means that the participants overall expressed biased ratings. The mean EEB was 0.45 (SD = 0.38), the mean EAB was 0.36 (SD = 0.55), and the mean SES was 0.53 (SD = 0.58) (Figure 1).

The participants’ means and standard deviations for the subscales of the German version of the Interpersonal Reactivity Index (Davis, 1983; Paulus, 2009) were in the expected range based on the data by Paulus (29.07.2024; https://www.cpaulus.de/tt-empathie.html). The mean for EC was 15.42 (SD = 2.17), for FS was 14.88 (SD = 3.07), for PD was 11.17 (SD = 2.68), and for PT was 15.67 (SD = 2.08).

### Imaging results

#### Rating phase: ROI analyses & whole brain analysis

Our ROI analysis for the rSMG (6mm sphere around 65 -37 33) yielded significant results for SOD (M = 1.06 (SD = 1.58), t(23) = 3.28, p < .01) and for SOD_other_ (M = 0.65 (SD = 1.33), t(23) = 2.38, p < .05) (see Fig. 2B). The rSMG did not show differential activity for the SOD_self_ contrast and the self-salience contrast (p > .10). However, the SOD_self_ and SOD_other_ contrasts did not differ significantly either (p > .10).

**Figure 2.**
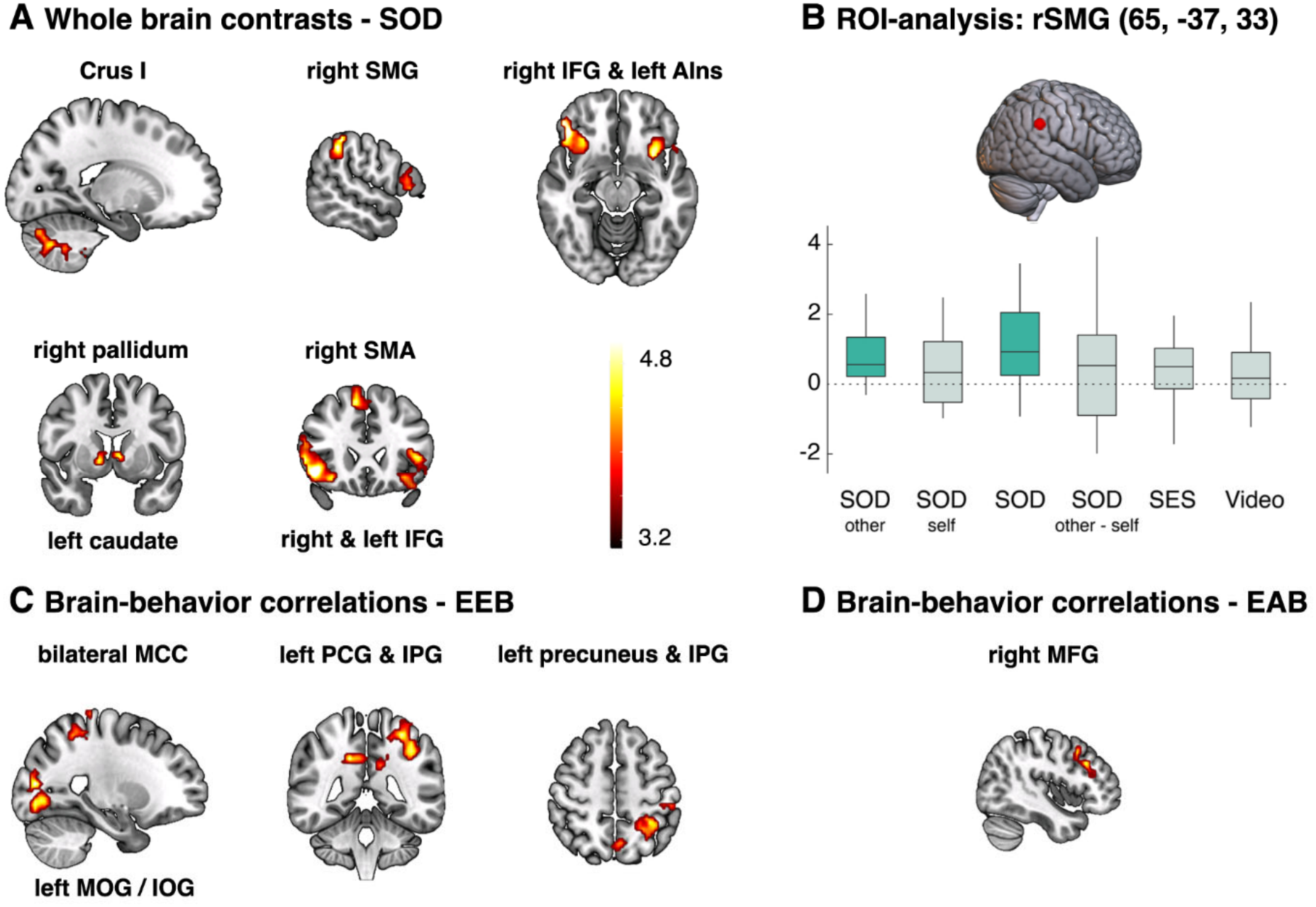
Imaging results for SOD, EEB, and EAB. T-values represented via color scale with higher values in yellow. **(A)** Whole brain contrast for incongruent vs. congruent trials. **(B)** Boxplots for differences in activity in the rSMG between incongruent and congruent trials. Significant differences are in dark green and insignificant differences are in light green. **(C)** Areas where activity correlated with the participants’ EEB. **(D)** Areas where activity correlated with participants’ EAB. Abbreviations: IFG = inferior frontal gyrus, Ins = Insular, IOG = inferior occipital gyrus, IPG = inferior parietal gyrus, MCC = middle cingulate cortex, MFG = middle frontal gyrus, MOG = middle occipital gyrus, PCG = postcentral gyrus, SMA = supplementary motor area, (r)SMG = (right) supramarginal gyrus.

In the whole brain analysis, we found significant differences in neural activity for the SOD, SOD_other_, and SOD_self_, and self-salience contrasts (see Table 1 and Fig. 2A & Fig. 3A). The SOD contrast was related to activity in the left anterior insula and right SMG among other regions. Again, there were no significant differences when contrasting the congruence effect in self- and other-related trials (SOD_self_ - SOD_other_). The self-salience contrast was related to activity in the bilateral PFC, left precuneus, left mid cingulate cortex (MCC), and rSMG. The clusters related to self-salience overlapped with those related to SOD in the rSMG, the rSFG, and the left IFG/Ins (see Fig. 3B).

**Table 1.** Rating phase: Results of whole brain analysis.

|  | Coordinates (mm) | Peak t-value | Cluster size | Cluster p-value (corr.) |
|---|---|---|---|---|
| **Incongruent > Congruent** |  |  |  |  |
| **SOD<sub>other</sub>; k<sub>critical</sub> = 87** |  |  |  |  |
| Left IFG (orb) & | **-27, 26, -13** | **6.24** | **465** | **<.001** |
| right pallidum | 12, 2, -1 | 5.60 |  |  |
| left caudate | -6, 2, 2 | 4.92 |  |  |
| Right IFG (orb & tri) | **51, 38, -10** | **5.75** | **102** | **.021** |
|  | 48, 20, -7 | 5.15 |  |  |
|  | 54, 26, -1 | 4.77 |  |  |
| Left IFG (orb & oper) | **-48, 20, -1** | **5.02** | **87** | **.037** |
|  | -57, 14, 8 | 4.89 |  |  |
| Right SMA | **9, 23, 50** | **5.00** | **106** | **.018** |
|  | 0, 14, 68 | 4.27 |  |  |
| **SOD<sub>self</sub>; k<sub>critical</sub> = 361** |  |  |  |  |
| Right IFG (orb & oper) & Ins<sup>1</sup> | **45, 29, -10** | **5.92** | **361** | **<.001** |
|  | 30, 17, -13, | 4.93 |  |  |
|  | 51, 17, 5 | 3.98 |  |  |
| **SOD; k<sub>critical</sub> = 75** |  |  |  |  |
| Right IFG (orb) | **45, 32, -13** | **6.38** | **452** | **<.001** |
|  | 51, 38, -10 | 6.31 |  |  |
|  | 45, 23, -7 | 5.84 |  |  |
| rSMG | **57, -40, 41** | **5.18** | **97** | **.020** |
|  | 60, -46, 26 | 3.58 |  |  |
| Left Ins<sup>1</sup> & IFG (orb) | **-30, 11, -13** | **5.10** | **263** | **<.001** |
|  | -36, 20, -19 | 4.94 |  |  |
|  | -48, 17, -7 | 4.88 |  |  |
| Right SMA & SFG (medial) | **9, 23, 53** | **4.93** | **91** | **.025** |
|  | 0, 26, 53 | 3.84 |  |  |
| Left Caudate & Right Pallidum | **-6, 5, 2** | 4.93 | 75 | .048 |
|  | 12, 5, -1 | 4.80 |  |  |
| Left cerebellum 7b, Crus1, & Lobule IX | **-12, -70, -40** | **4.75** | **212** | **<.001** |
|  | -18, -73, -34 | 4.43 |  |  |
|  | -3, -55, -40 | 4.32 |  |  |
| **Self > Other** |  |  |  |  |
| **Self-salience; k<sub>critical</sub> = 83** |  |  |  |  |
| Right SFG & MFG (DLPFC) | **24, 50, 32** | **4.79** | **246** | **<.001** |
|  | 30, 53, 14 | 4.76 |  |  |
|  | 24, 47, 23 | 4.36 |  |  |
| Left precuneus & left MCC | **-3, -67, 50** | **4.61** | **1304** | **<.001** |
|  | -6, -31, 32 | 4.50 |  |  |
|  | -12, -55, 29 | 4.39 |  |  |
| Left SFG & MFG (DLPFC) | **-27, 59, 14** | **4.54** | **196** | **.001** |
|  | -39, 50, 14 | 4.53 |  |  |
|  | -42, 41, 23 | 3.90 |  |  |
| Right SMG | **51, -46, 32** | **5.75** | **107** | **.019** |
| Left SFG & MFG | **-18, 11, 53** | **5.52** | **175** | **.002** |
|  | -27, 5, 65 | 4.43 |  |  |
|  | -33, 5, 53 | 4.34 |  |  |
| Right SMA & SFG | **9, 23, 65** | **5.41** | **91** | **.034** |
|  | 18, 14, 62 | 4.17 |  |  |
|  | 15, 29, 50 | 3.99 |  |  |
| Left IFG (orb) & left Ins | **-27, 32, -10** | **5.29** | **117** | **.013** |
|  | -33, 23, -22 | 4.15 |  |  |
|  | -27, 12, -13 | 4.04 |  |  |
| Left caudate | **-12, -1, 17** | **4.72** | **83** | **.045** |
|  | -9, 8, 11 | 3.90 |  |  |

*Notes.* Results are presented in Montreal Neurological Institute coordinate system (MNI). Peak coordinates were localized using the AAL3 (Rolls et al., 2020). List of abbreviations: DLPFC = dorsolateral prefrontal cortex, IFG = inferior frontal gyrus, Ins = insula, MFG = middle frontal gyrus, orb = pars orbitalis, oper = pars opercularis, rSMG = right supramarginal gyrus, SFG = superior frontal gyrus, SMA = supplementary motor area, tri = pars triangularis.
<sup>1</sup> Anterior Insula according to Neuromorphometrics.

**Figure 3.**
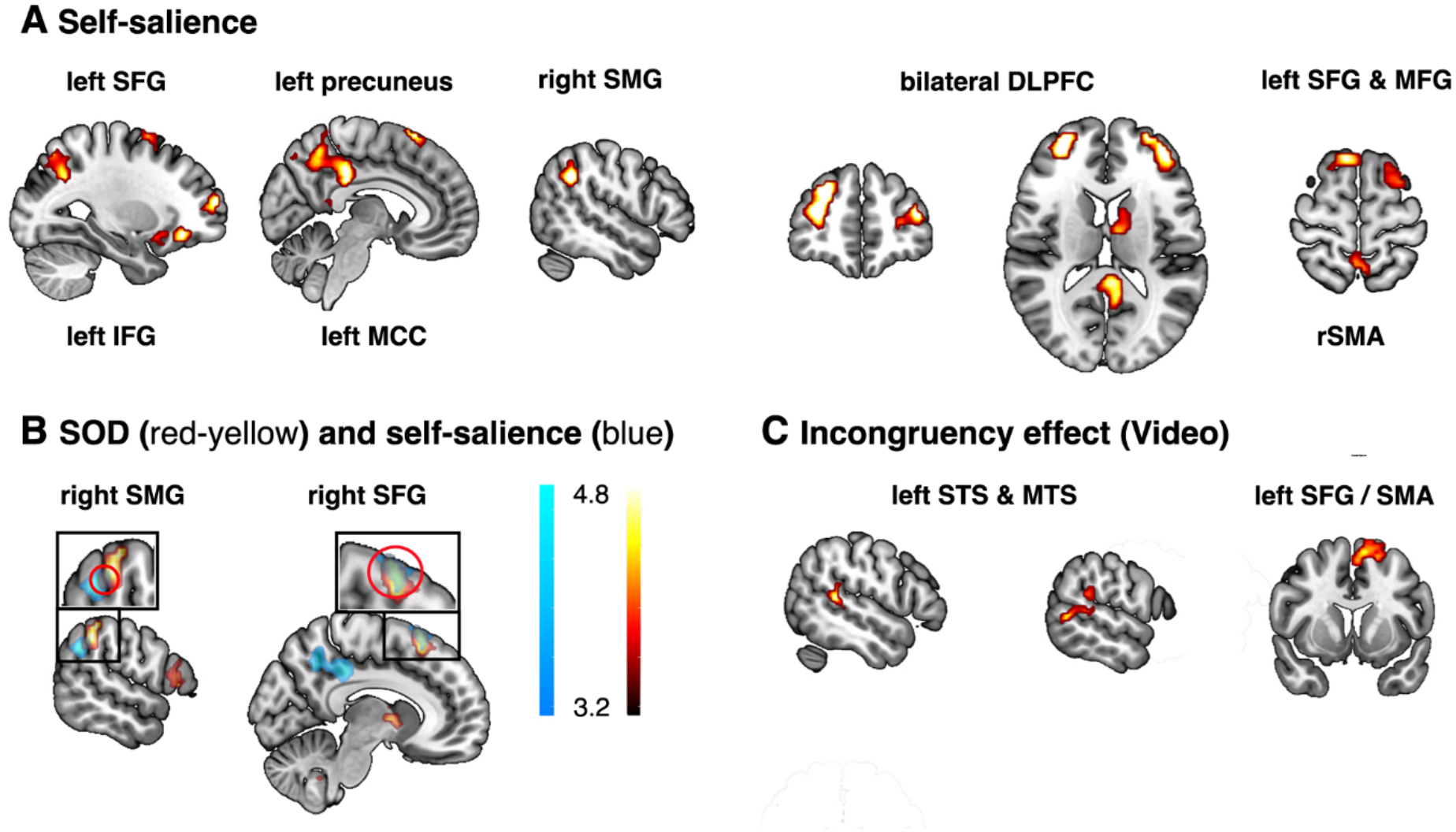
Imaging results for self-salience and the video presentation. T-values represented via color scales with higher values in yellow/light blue. **(A)** Whole brain contrast for congruent self vs. congruent other trials. **(B)** Overlay (red circle) of activity related to SOD and self-salience. Zoomed in on the right SMG (left) and right SFG (right). **(C)** Whole brain contrast for the presentation of incongruent vs. congruent videos. Abbreviations: DLPFC = dorsolateral prefrontal cortex, IFG = inferior frontal gyrus, MCC = middle cingulate cortex, MFG = middle frontal gyrus, MTS = middle temporal sulcus, SFG = superior frontal gyrus, SMA = supplementary motor area, (r)SMG = (right) supramarginal gyrus, STS = superior temporal sulcus.

#### Brain-behavior correlations

We used multiple regressions to test whether interindividual variability in participants’ biases was related to changes in neural activity. Our analysis revealed that the strength of the EEB was negatively correlated with the congruency effect in the left inferior parietal gyrus, the left inferior occipital gyrus, precuneus, as well as the MCC (*k*_*critical*_ > 241, see Fig. 2C). The strength of the EAB was negatively correlated with the congruency effect in the right DLPFC (*k*_*critical*_ > 88; s. Table 2 and Fig. 2D). For the SES we could not find any significant brain-behavior correlations.

**Table 2.** Rating phase: Results of multiple regressions for EEB & EAB.

|  | Coordinates (mm) | Peak t-value | Cluster size | Cluster p-value (corr.) |
|---|---|---|---|---|
| **EEB(neg) vs SOD<sub>other</sub>; k<sub>critical</sub> = 241** |  |  |  |  |
| Left MOG & IOG | **-21, -82, 11** | **5.61** | **317** | **<.001** |
|  | -42, -73, -4 | 5.10 |  |  |
|  | -27, -76, -1 | 5.03 |  |  |
| Left IPG & PCG | **-39, -46, 41** | **5.46** | **328** | **<.001** |
|  | -30, -43, 56 | 5.01 |  |  |
|  | -24, -49, 53 | 4.37 |  |  |
| Bilateral MCC & left precuneus | **15, -43, 35** | **4.09** | **241** | **<.001** |
|  | -12, -43, 29 | 3.62 |  |  |
|  | -6, -61, 53 | 3.61 |  |  |
| **EAB(neg) vs SOD<sub>self</sub>; k<sub>critical</sub> = 88** |  |  |  |  |
| Right MFG (DLPFC) | **45, 23, 32** | **5.15** | **88** | **.020** |
|  | 42, 14, 41 | 4.67 |  |  |
|  | 42, 29, 17 | 4.16 |  |  |

*Notes.* Results are presented in MNI. Peak coordinates were localized using the AAL3 (Rolls et al., 2020). List of abbreviations: DLPFC = dorsolateral prefrontal cortex, IOG = inferior occipital gyrus, IPG = inferior parietal gyrus, MCC = mid cingulate gyrus, MFG = middle frontal gyrus, MOG = middle occipital gyrus, PCG = postcentral gyrus.

#### Incongruence effect during video presentation

For this exploratory analysis, we replaced the regressor for video with four regressors specifying the condition of video presentation (e.g., happy video with congruent picture). During video presentation, incongruency led to higher activity in the lSMA, left superior frontal gyrus, and left middle and superior temporal gyrus (see Table 3 and Fig. 3).

**Table 3.** Video presentation phase: Results of whole-brain analysis.

|  | **Coordinates (mm)** | **Peak t-value** | **Cluster size** | **Cluster p-value (corr.)** |
|---|---|---|---|---|
| **Incongruent > congruent (all trials)** |  |  |  |  |
| **k<sub>critical</sub> = 120** |  |  |  |  |
| Left STG & MTG | **-51, -40, 8** | **4.94** | **120** | **.023** |
|  | -57, -55, 2 | 4.58 |  |  |
|  | -60, -34, 11 | 4.46 |  |  |
| Left SFG & SMA | **-9, 8, 62** | **4.83** | **122** | **.022** |
|  | -3, 8, 56 | 4.40 |  |  |
|  | -18, 8, 59 | 4.36 |  |  |

*Notes.* Results are presented in MNI. Peak coordinates were localized using the AAL3 (Rolls et al., 2020). List of abbreviations: MTG = middle temporal gyrus, SFG = superior frontal gyrus, SMA = supplementary motor area, STG = superior temporal gyrus

## Discussion

In this fMRI study, we investigated the neural processes involved in affective self-other distinction. We employed a novel paradigm that presented participants with food stimuli eliciting happiness or disgust and short videos of another person’s happy or disgusted facial expression (Goregliad Fjaellingsdal et al., 2025). When participants had to evaluate their own or another person’s emotion in situations of incongruence, the rSMG and anterior insula among other regions were activated. Interindividual variability in the egocentricity bias (EEB) and the altercentricity bias (EAB) correlated with distinct brain areas, suggesting partly dissociable neural processes. Finally, when participants evaluated their own emotions in comparison to others’ congruent emotions, we observed activations in parts of the so-called default network, as well as in the rSMG. This suggests that the rSMG subserves processes related to both self-salience and self-other distinction, emphasizing the interrelation between these constructs.

### Behavioral results

As expected, participants rated emotions as less intense in the incongruent conditions than in the congruent conditions. This was true when rating their own emotions and the emotions of others, reflecting an altercentric and egocentric bias, respectively. Additionally, participants exhibited a self-salience bias, rating their own emotions as generally more intense than others’.

The behavioral results were generally consistent with those of our previous studies using the same paradigm (Goregliad Fjaellingsdal et al., 2025). However, unlike these previous behavioral studies, the EEB and EAB did not differ in the current fMRI study. Previously (Goregliad Fjaellingsdal et al., 2025), we showed that the EAB is more state-dependent and variable than the EEB and self-salience, so situational factors in the current study, most notably the MRI setting, may have increased the EAB. This finding is consistent with previous studies on affective SOD using different paradigms (Bukowski et al., 2020; Silani et al., 2013). Future studies should address which situational factors influence the EAB.

### The role of the rSMG in affective self-other distinction

As hypothesized, we observed higher activity in the rSMG when the participants experienced incongruent emotional states and had to evaluate either their own or the other person’s emotional state. This emphasizes the role of the rSMG in affective SOD suggested by previous work (Bukowski et al., 2020; Silani et al., 2013; Steinbeis et al., 2015). In the FEEB, participants have to assess the other person’s emotional state directly based on facial cues, unlike in previous paradigms which asked for an indirect inference based on the context (Silani et al., 2013; Steinbeis et al., 2015; von Mohr et al., 2019). Evaluating the other’s emotions in the FEEB might therefore rely more on the experience sharing route, whereas it relies on the mentalizing or theory-of-mind route in previous SOD paradigms. Our results show that the rSMG responds to situations of affective incongruence, independently of the route through which the other person’s emotions were experienced. Our results also show that the rSMG has a supramodal function in resolving emotional incongruency (Quesque & Brass, 2019). It is involved not only in processing emotional states elicited via visuo-tactile stimulation (Bukowski et al., 2020; Riva et al., 2022; Silani et al., 2013) and monetary gains and losses (Steinbeis et al., 2015), but also via visuo-gustatory stimuli. Finally, our results replicate the observation that the rSMG is related to SOD in general and not specifically in other-judgment trials (Bukowski et al., 2020). This shows that the rSMG is activated by situations calling for affective self-other distinction, irrespective of whether the focus is on the self or the other.

### Beyond the rSMG: Neural networks involved in affective SOD

The AIns and ACC are strongly implicated in experience sharing (Bernhardt & Singer, 2012; Engen & Singer, 2013) and the salience network (Seeley, 2019). Furthermore, the AIns is also involved in affective processing and processing gustatory stimuli (Lamm & Singer, 2010; Uddin et al., 2017). Therefore, we expected higher activity in the AIns and ACC during states of emotional incongruency. Indeed, we observed higher activity in the AIns, replicating other studies on affective SOD (Bukowski et al., 2020; Steinbeis et al., 2015). However, contrary to our hypothesis and to another study on SOD (Steinbeis et al., 2015), we did not find higher activity in the ACC. The AIns and ACC may serve distinct roles in the salience network (Seeley, 2019) and in processing sensory vs. motivational and behavioral aspects of emotions (Bernhardt & Singer, 2012; Craig, 2002, 2009). It is possible that the incongruency in emotional states is resolved rather on the level of sensory representations than on the level of behavioral output. Our results support the notion that the AIns is involved in processing emotional incongruency/SOD regardless of the modality (Bukowski et al., 2020).

### Brain-behavior correlations

As our final sample comprised only 24 participants, the analyses of brain-behavior correlations were underpowered and should be considered with caution. Notably, the EEB correlated negatively with activity in areas related to low level facial perceptions (lOG; de Haas et al., 2021), the perception of dynamic facial expressions (inferior parietal gyrus; Sarkheil et al., 2013) and to activity in the precuneus and the MCC/PCC (Leech & Smallwood, 2019). Interindividual variability in the EAB on the other hand was negatively related to activity in the right dlPFC. The strength of the left dlPFC - rSMG connectivity has been linked to a smaller EEB (Steinbeis et al., 2015), and Bukowski and colleagues (2020) therefore argued that at least in some individuals SOD was related to cognitive control. Interestingly however, neither EEB nor EAB was correlated with the rSMG, which suggests that rSMG activity cannot explain interindividual variability. Moreover, the brain-behavior correlations for EEB and EAB were different, which suggests that they rely on overlapping, but at least partly dissociable neural processes.

### Experiencing emotional incongruency vs. resolving emotional incongruency

Our trial structure allowed us to disentangle the processes elicited by the incongruence itself (video phase; Figure 1) and the processes involved in resolving this incongruence (rating phase). When confronted with emotionally incongruent stimuli, we found higher activity around the posterior parts of the superior temporal sulcus which has been linked to perception of biological movements, face processing, and theory of mind (for an overview see Hein & Knight, 2008). Importantly, however, we did not find higher activity in the rSMG during incongruent videos. This is in contrast to another fMRI study on affective SOD, which also found a congruency effect on rSMG activity in non-judgement trials (Steinbeis et al., 2015). Our results suggest that the rSMG is not activated when simply experiencing emotional incongruency, but only when having to resolve the incongruency to rate either one’s own or the other’s emotion. The difference between studies might be due to different emotional stimuli and task demands. We would like to emphasize, though, that the FEEB paradigm is helpful in distinguishing these different processes involved in affective SOD.

### Self-salience and affective self-other distinction

Previous imaging studies on affective SOD have not distinguished the neural correlates of SOD and self-salience. As the default mode network (DMN) reflects self-referential processes (Raichle, 2015; Raichle et al., 2001), we expected, and found, higher activity in DMN regions during self-congruent compared to other-congruent judgements (i.e., precuneus and MCC/PCC; Raichle, 2015; Smallwood et al., 2021). Interestingly, we observed that higher activity in the rSMG and the right SMA/SFG was found for both the self-salience contrast and the SOD contrast (Fig. 3). The rSMG area activated for self-salience was overlapping with the rSMG area activated in SOD but extended further posterior (see Fig. 3B). Based on our finding of rSMG’s involvement in both SOD and self-salience, we suggest that the rSMG is relevant for representing the self in contrast to others, not only in situations of incongruency. Support comes from research on interoception, which shows that interoceptive and exteroceptive signals of the bodily state converge in the SMG (Salvato et al., 2020). Better interoceptive self-awareness might facilitate SOD under high demands (Palmer & Tsakiris, 2018), which might explain our previous behavioral finding of a negative correlation between the self-salience effect and the EEB (Goregliad Fjaellingsdal et al., 2025).

### Limitations

Due to our strict exclusion criteria, our final sample consisted of only 24 participants. We therefore did not have enough power to reliably detect brain behavior correlations. Since the data acquisition of this fMRI study, we developed an adapted version of the FEEB with an extended prerating (Goregliad Fjaellingsdal et al., 2025). Future studies with the FEEB should thus use this adapted version to prevent having to exclude too many participants.

## Conclusions

We could show that emotional incongruency elicited via experience sharing activated the rSMG and the AIns, which expands on previous studies that elicited emotional incongruency via mentalizing. Additionally, we could show that self-salience is related to activity in areas of the DMN and the rSMG. Our study shows that the rSMG is involved in representing self-related information in situations of self-other distinction also independently of incongruency between own and other’s emotional state. Future research should further explore the contribution of interoception and self-awareness to SOD.

## Acknowledgements

This work was supported by the Deutsche Forschungsgemeinschaft (DFG, German Research Foundation) – CRC-1665-515637292.

## References

Bernhardt, B. C., & Singer, T. (2012). The neural basis of empathy. Annu Rev Neurosci, 35, 1–23. 10.1146/annurev-neuro-062111-150536

Blechert, J., Lender, A., Polk, S., Busch, N. A., & Ohla, K. (2019). Food-Pics_Extended—An Image Database for Experimental Research on Eating and Appetite: Additional Images, Normative Ratings and an Updated Review [Original Research]. Front Psychol, Volume 10 - 2019. 10.3389/fpsyg.2019.00307

Brainard, D. H. (1997). The Psychophysics Toolbox. Spatial Vision, 10(4), 433–436. 10.1163/156856897X00357

Bukowski, H., Tik, M., Silani, G., Ruff, C. C., Windischberger, C., & Lamm, C. (2020). When differences matter: rTMS/fMRI reveals how differences in dispositional empathy translate to distinct neural underpinnings of self-other distinction in empathy. Cortex, 128, 143–161. 10.1016/j.cortex.2020.03.009

Bukowski, H., Todorova, B., Boch, M., Silani, G., & Lamm, C. (2021). Socio-cognitive training impacts emotional and perceptual self-salience but not self-other distinction. Acta Psychol (Amst), 216, 103297. 10.1016/j.actpsy.2021.103297

Ciocca, G., Napoletano, P., & Schettini, R. (2018). CNN-based features for retrieval and classification of food images. Computer Vision and Image Understanding, 176-177, 70–77. 10.1016/j.cviu.2018.09.001

Craig, A. D. (2002). How do you feel? Interoception: the sense of the physiological condition of the body. Nature Reviews Neuroscience, 3(8), 655–666. 10.1038/nrn894

Craig, A. D. (2009). How do you feel--now? The anterior insula and human awareness. Nature Reviews Neuroscience, 10(1), 59–70. 10.1038/nrn2555

Davis, M. H. (1983). Measuring individual differences in empathy: Evidence for a multidimensional approach Journal of Personality and Social Psychology, 44(1), 113–126.

de Guzman, M., Bird, G., Banissy, M. J., & Catmur, C. (2016). Self-other control processes in social cognition: from imitation to empathy. Philos Trans R Soc Lond B Biol Sci, 371(1686), 20150079. 10.1098/rstb.2015.0079

de Haas, B., Sereno, M. I., & Schwarzkopf, D. S. (2021). Inferior Occipital Gyrus Is Organized along Common Gradients of Spatial and Face-Part Selectivity. J Neurosci, 41(25), 5511–5521. 10.1523/JNEUROSCI.2415-20.2021

De Meulemeester, C., Lowyck, B., & Luyten, P. (2021). The role of impairments in self-other distinction in borderline personality disorder: A narrative review of recent evidence. Neurosci Biobehav Rev, 127, 242–254. 10.1016/j.neubiorev.2021.04.022

Eddy, C. M. (2018). Social cognition and self-other distinctions in neuropsychiatry: Insights from schizophrenia and Tourette syndrome. Prog Neuropsychopharmacol Biol Psychiatry, 82, 69–85. 10.1016/j.pnpbp.2017.11.026

Engen, H. G., & Singer, T. (2013). Empathy circuits. Current Opinion in Neurobiology, 23(2), 275–282. 10.1016/j.conb.2012.11.003

Frith, C. D., & Frith, U. (2006). The neural basis of mentalizing. Neuron, 50(4), 531–534. 10.1016/j.neuron.2006.05.001

Goregliad Fjaellingsdal, T., Dohr, K., Zschoche, G., & Kramer, U. M. (2025). Advancing the measurement of affective self-other distinction: A novel paradigm to measure intra- and inter-individual differences in emotional biases. Acta Psychol (Amst), 261, 105840. 10.1016/j.actpsy.2025.105840

Hartmann, H., Lengersdorff, L., Hitz, H. H., Stepnicka, P., & Silani, G. (2022). Emotional Ego- and Altercentric Biases in High-Functioning Autism Spectrum Disorder: Behavioral and Neurophysiological Evidence. Front Psychiatry, 13, 813969. 10.3389/fpsyt.2022.813969

Hein, G., & Knight, R. T. (2008). Superior Temporal Sulcus-It’s My Area: Or Is It? J Cogn Neurosci, 20(12), 2125–2136.

Janke, S., & Glöckner-Rist, A. (2012). Deutsche Version der Positive and Negative Affect Schedule (PANAS). Zusammenstellung Sozialwissenschaftlicher Items und Skalen (ZIS). 10.6102/zis146

Kleiner, M., Brainard, D. H., Pelli, D., Ingling, A., Murray, R., & Broussard, C. (2007). What’s new in Psychtoolbox-3. Perception, 36, 1–16. 10.1068/v070821

Lamm, C., Bukowski, H., & Silani, G. (2016). From shared to distinct self-other representations in empathy: evidence from neurotypical function and socio-cognitive disorders. Philos Trans R Soc Lond B Biol Sci, 371(1686), 20150083. 10.1098/rstb.2015.0083

Lamm, C., & Singer, T. (2010). The role of anterior insular cortex in social emotions. Brain Struct Funct, 214(5-6), 579–591. 10.1007/s00429-010-0251-3

Leech, R., & Smallwood, J. (2019). The posterior cingulate cortex: Insights from structure and function. Handb Clin Neurol, 166, 73–85. 10.1016/B978-0-444-64196-0.00005-4

wLieberman, M. D. (2007). Social cognitive neuroscience: a review of core processes. Annu Rev Psychol, 58, 259–289. 10.1146/annurev.psych.58.110405.085654

Mars, R. B., Neubert, F. X., Noonan, M. P., Sallet, J., Toni, I., & Rushworth, M. F. (2012). On the relationship between the “default mode network” and the “social brain”. Front Hum Neurosci, 6, 189. 10.3389/fnhum.2012.00189

Palmer, C. E., & Tsakiris, M. (2018). Going at the heart of social cognition: is there a role for interoception in self-other distinction? Current Opinion in Psychology, 24, 21–26. 10.1016/j.copsyc.2018.04.008

Paulus, C. (2009). DER SAARBRÜCKER PERSÖNLICHKEITSFRAGEBOGEN SPF(IRI) ZUR MESSUNG VON EMPATHIE: Psychometrische Evaluation der deutschen Version des Interpersonal Reactivity Index. 10.23668/psycharchives.9249

Pelli, D. G. The VideoToolbox software for visual psychophysics: transforming numbers into movies. (0169-1015 (Print)).

Quesque, F., & Brass, M. (2019). The Role of the Temporoparietal Junction in Self-Other Distinction. Brain Topography, 32(6), 943–955. 10.1007/s10548-019-00737-5

Raichle, M. E. (2015). The brain’s default mode network. Annu Rev Neurosci, 38, 433–447. 10.1146/annurev-neuro-071013-014030

Raichle, M. E., MacLeod, A. M., Snyder, A. Z., Powers, W. J., Gusnard, D. A., & Shulman, G. L. (2001). A default mode of brain function. Proc Natl Acad Sci U S A, 98(2), 676–682. 10.1073/pnas.98.2.676

Riva, F., Lenger, M., Kronbichler, M., Lamm, C., & Silani, G. (2022). The role of right supra-marginal gyrus and secondary somatosensory cortex in age-related differences in human emotional egocentricity. Neurobiol Aging, 112, 102–110. 10.1016/j.neurobiolaging.2022.01.002

Riva, F., Triscoli, C., Lamm, C., Carnaghi, A., & Silani, G. (2016). Emotional Egocentricity Bias Across the Life-Span. Front Aging Neurosci, 8, 74. 10.3389/fnagi.2016.00074

Rolls, E. T., Huang, C. C., Lin, C. P., Feng, J., & Joliot, M. (2020). Automated anatomical labelling atlas 3. Neuroimage, 206, 116189. 10.1016/j.neuroimage.2019.116189

Salvato, G., Richter, F., Sedeno, L., Bottini, G., & Paulesu, E. (2020). Building the bodily self-awareness: Evidence for the convergence between interoceptive and exteroceptive information in a multilevel kernel density analysis study. Hum Brain Mapp, 41(2), 401–418. 10.1002/hbm.24810

Sarkheil, P., Goebel, R., Schneider, F., & Mathiak, K. (2013). Emotion unfolded by motion: a role for parietal lobe in decoding dynamic facial expressions. Soc Cogn Affect Neurosci, 8(8), 950–957. 10.1093/scan/nss092

Schurz, M., Maliske, L., & Kanske, P. (2020). Cross-network interactions in social cognition: A review of findings on task related brain activation and connectivity. Cortex, 130, 142–157. 10.1016/j.cortex.2020.05.006

Seeley, W. W. (2019). The Salience Network: A Neural System for Perceiving and Responding to Homeostatic Demands. J Neurosci, 39(50), 9878–9882. 10.1523/JNEUROSCI.1138-17.2019

Silani, G., Lamm, C., Ruff, C. C., & Singer, T. (2013). Right supramarginal gyrus is crucial to overcome emotional egocentricity bias in social judgments. J Neurosci, 33(39), 15466–15476. 10.1523/JNEUROSCI.1488-13.2013

Smallwood, J., Bernhardt, B. C., Leech, R., Bzdok, D., Jefferies, E., & Margulies, D. S. (2021). The default mode network in cognition: a topographical perspective. Nature Reviews Neuroscience, 22(8), 503–513. 10.1038/s41583-021-00474-4

Steinbeis, N. (2016). The role of self-other distinction in understanding others’ mental and emotional states: neurocognitive mechanisms in children and adults. Philos Trans R Soc Lond B Biol Sci, 371(1686), 20150074. 10.1098/rstb.2015.0074

Steinbeis, N., Bernhardt, B. C., & Singer, T. (2015). Age-related differences in function and structure of rSMG and reduced functional connectivity with DLPFC explains heightened emotional egocentricity bias in childhood. Soc Cogn Affect Neurosci, 10(2), 302–310. 10.1093/scan/nsu057

Uddin, L. Q., Nomi, J. S., Hebert-Seropian, B., Ghaziri, J., & Boucher, O. (2017). Structure and Function of the Human Insula. J Clin Neurophysiol, 34(4), 300–306. 10.1097/WNP.0000000000000377

von Mohr, M., Finotti, G., Ambroziak, K. B., & Tsakiris, M. (2019). Do you hear what I see? An audiovisual paradigm to assess emotional egocentricity bias. Cogn Emot, 1–15. 10.1080/02699931.2019.1683516

Yitzhak, N., Giladi, N., Gurevich, T., Messinger, D. S., Prince, E. B., Martin, K., & Aviezer, H. (2017). Gently does it: Humans outperform a software classifier in recognizing subtle, nonstereotypical facial expressions. Emotion, 17(8), 1187–1198. 10.1037/emo0000287

