## supplementary materials for "Resolving the neural processes involved in affective self-other distinction – an fMRI study"

### Supplementary Material

Tatiana Goregliad Fjaellingsdal[1,2,‡], Konstantin Döhr[3,‡], Emily Fitzgibbon[1], Martin Göttlich[3], Ulrike M. Krämer[3]

**Table S1**

*Mean and standard deviations for PANAS scores*

| | NA | PA |
|---|---|---|
| **Pre FEEB** | 13.27 (2.84) | 31.51 (6.12) |
| **Post FEEB** | 12.04 (2.10) | 30.18 (6.52) |

*Note.* Cells contain mean values with (SD in brackets), NA = negative affect subscale, PA = positive affect subscale, PANAS = Positive Affect and Negative Affect Schedule.

**Table S2**

*Spearman rank correlations between bias scores and questionnaire subscales*

| **Measures** | **IRI** | | | | **PANAS** | | | |
|---|---|---|---|---|---|---|---|---|
| | | | | | **Pre FEEB** | | **Post FEEB** | |
| **Subscales** | **EC** | **FS** | **PD** | **PT** | **NA** | **PA** | **NA** | **PA** |
| **EEB** | .13 | .30 | **.44*** | -.04 | .06 | -.05 | -.16 | .02 |
| **EAB** | .07 | .25 | .18 | .07 | -.03 | -.32 | .02 | -.30 |
| **SES** | **.50*** | -.00 | .33 | .11 | -.03 | .09 | -.02 | .03 |

*Note.* Cells contain spearman rank correlations (rho), EC = empathic concern subscale, FS = fantasy subscale, IRI = Interpersonal Reactivity Index, NA = negative affect subscale, PA = positive affect subscale, PANAS = Positive Affect and Negative Affect Schedule, PD = personal distress subscale, PT = perspective taking subscale.

* $p < .05$.
